# A Novel Adhesin of *B. pertussis* is Key to Colonisation of Epithelial Cells

**DOI:** 10.1101/2024.09.23.613229

**Authors:** Michael Gollan, Monica C Gestal, Katelyn M Parrish, Eric T Harvill, Andrew Preston, Iain MacArthur

## Abstract

Despite effective vaccines to protect against Whooping cough, or pertussis, the disease is resurgent in many countries. A switch from acellular to whole-cell vaccines has resulted in waning protective immunity, likely contributing to increases in infection prevalence, underlining the need to better understand *B. pertussis* virulence. As a respiratory pathogen, *B. pertussis* colonises the upper respiratory tract utilising an array of adhesins, four of which (FHA, pertactin, Fim2/3) are included in the acellular vaccine. In this study, we identified two Bvg regulated genes that are upregulated during virulence conditions and thus potentially involved in pathogenesis. *bp1251* and *bp1252* encode orphan toxin B subunits, with homology to AB toxin B subunits. Mutation of *bp1251* and *bp1252* reduced the *in vitro* adherence of *B. pertussis* to A549 and BEAS-2B alveolar and bronchial epithelial-like cells. In a murine model of infection, *bp1251* and *bp*1252 mutant strains were recovered from the nasal cavity and lungs at lower levels than WT. *In vitro* no effect of mutation of *bp1251* or *bp1252* on cell invasion or toxicity was found suggesting that these proteins do not form part of a toxin. Given their homology to B subunits of AB toxins, and their role in colonisation, we propose that BP1251 and BP1252 are novel adhesins. Our data suggests that these proteins play a significant role in *Bordetella* infection and have the potential to further the understanding of *B. pertussis* pathogenesis.

## Introduction

Whooping Cough, also known as Pertussis, is a disease resulting from infection by the respiratory human pathogen *Bordetella pertussis*. Global vaccination schedules have largely controlled the incidence of Pertussis since the 1960s, when whole-cell vaccines (wP) were introduced [1,2]. Due to unfavourable reactogenicity, including suspected neurological side effects, a safer acellular vaccine (aP) was adopted in many high-income countries, with the wP vaccine retained in many low/middle-income countries. Although aP vaccines are very effective at preventing systemic disease, they have several limitations, including shorter-lasting protection [3–6], and inducing selection pressures for evolution of *B. pertussis* away from vaccine strains [7,8]. Furthermore, aP vaccination fails to prevent colonisation of the upper respiratory tract [9,10], potentially allowing high rates of carriage and transmission. Thus aP vaccines are considered to be less effective against currently circulating strains, phenomena seldom observed in countries using the wP vaccine [11–13][14–16]. These observations have highlighted the necessity to improve our understanding of *B. pertussis* pathogenesis.

*B. pertussis* virulence is regulated via the two-component *Bordetella virulence gene* (Bvg) system through which a sensory kinase (BvgS) relays external stimuli to a transcription factor (BvgA), thus modulating gene expression. Three Bvg states are currently accepted, representing virulent (Bvg+), avirulent (Bvg-), and intermediate (Bvgi) states. Most virulence factors, including the aP components filamentous haemagglutinin (FHA), pertussis toxin (PTx), pertactin (prn), and fimbriae (fim) are expressed in the Bvg+ state. The Bvgi state is characterised by high expression of adhesins and lower relative expression of other virulence factors [17,18] leading to suggestions that it is a transmission-focused state [19]. Colonisation is a vital first step for the bacterium to infect the host, with adherence to respiratory tissues required for the infection to proceed. [17]

We have characterised two co-transcribed genes, *bp1251* and *bp1252*, with ∼30% and 27% amino acid identity to pertussis toxin subunits PtxB/C and subtilase cytotoxin subunit B (SubB) of *Salmonella enterica*, respectively. These proteins have previously been identified as HotA and HotB (Homolog Of Toxin) [20], but have not been characterised. PtxB and C and SubB are B subunits of AB toxins, the components responsible for binding to the surface of target cells, but *bp1251* and *bp1252* do not appear to be part of a larger toxin-encoding locus. RNA-Seq showed that these “orphan” toxin B subunits are under BvgAS regulation and are transcribed in both Bvg+ and Bvgi states. It has been demonstrated that, in the absence of a cognate A toxin, B subunits can bind eukaryotic cells [21,22]. In this study we aimed to characterise these orphan B subunits and determine their importance to *B. pertussis* virulence.

## Materials and Methods

### Bacterial strains and culture conditions

Bacterial strains and plasmids used in this study are listed in Table 1 and Table 2 respectively*. B. pertussis* strains were cultured on charcoal agar (Scientific Laboratory Supplies). Strains were grown in Stainer-Scholte (SS) media [23,24] when liquid culture was required. The auxotrophic *E. coli* mutant ST18 was used as a conjugation donor [25].

**Table 1.** Strains used in this study.

| Strain | Description | Reference |
| --- | --- | --- |
| *B. pertussis* BP536 | WT | Mishra et al (2005) |
| *E. coli* ST18 | $\Delta hemA$ | Thoma et al (2009) |
| *E. coli* DH10b | Cloning strain | NEB |
| $\Delta BP1251$ | Deletion of *bp1251* | This study |
| $\Delta BP1252$ | Deletion of *bp1252* | This study |
| $\Delta BP1251$-1252 | Deletion of *bp1251* and<br>*bp1252* | This study |
| BP1251Comp | Complemented BP1251<br>deletion strain | This study |
| BP1252Comp | Complemented BP1252<br>deletion strain | This study |
| $\Delta FhaB$ | Deletion of *fhaB* | This study |
| $\Delta Fim$ | Deletion of *fimB-D* | This study |

**Table 2.** Plasmids used in this study.

| Plasmid | Description | Reference |
| --- | --- | --- |
| pSS4940(GG) | pSS4245-based suicide<br>plasmid (Gm<sup>R</sup>) for Golden<br>Gate cloning | MacArthur et al (2019) |
| pBBRKGG | pBBR-based shuttle vector<br>(Kan<sup>R</sup>) for Golden Gate<br>cloning | MacArthur et al (2019) |

### Molecular Biology

KO mutants were generated through homologous recombination. In-frame deletions were created by amplifying regions approximately 500bp upstream and downstream of the deletion. Primers used are listed in Table 3. These regions were cloned into pSS4940GG suicide vector using Golden Gate assembly [23]. Extrachromosomal complementation of the deletion was achieved through introducing the gene of interest under its native promoter carried on the plasmid pBBRK (Table 2). The pBBRK constructs were assembled through Golden Gate assembly and transformed into chemically competent *E. coli* ST18 cells.[22]

**Table 3.**
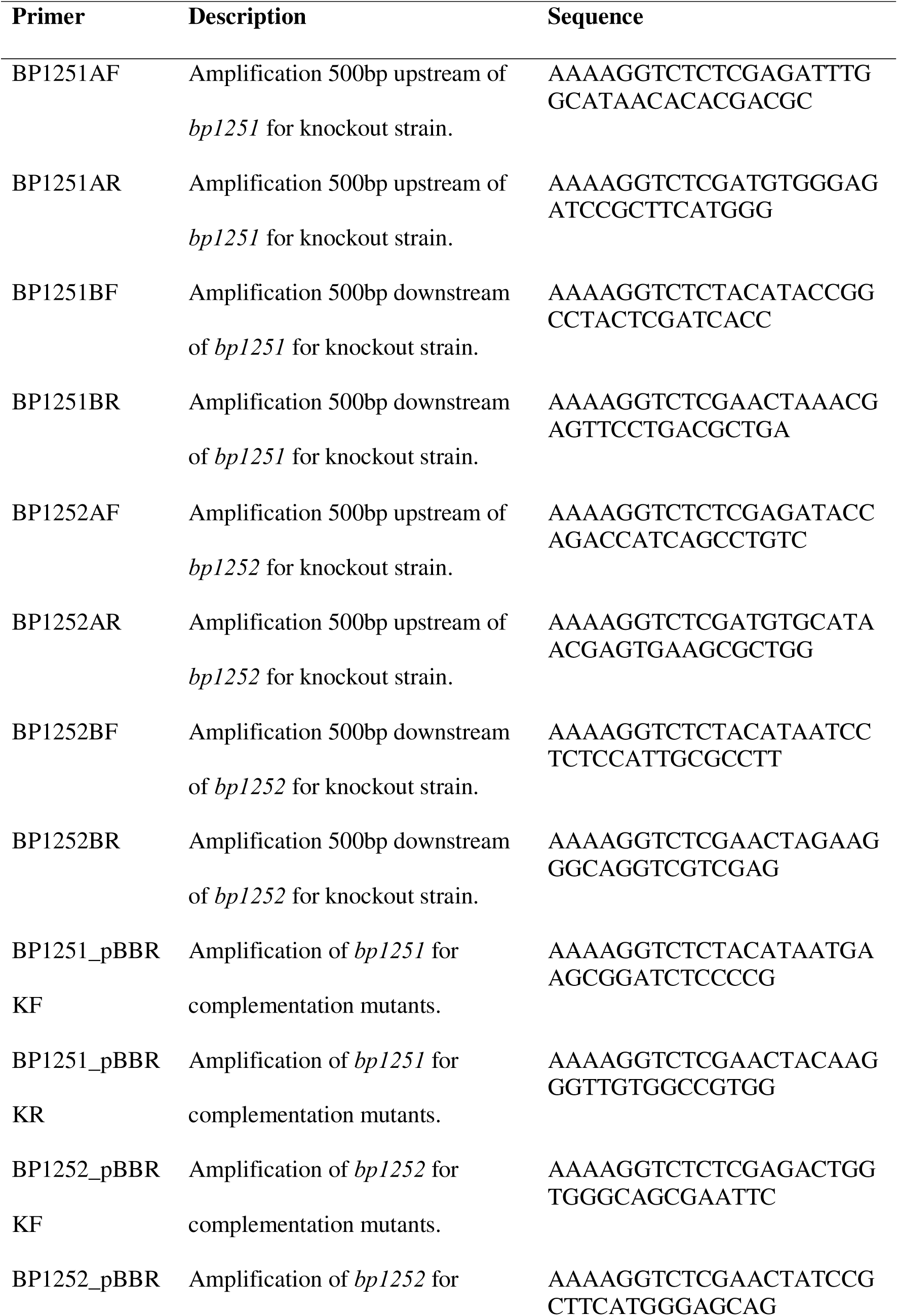
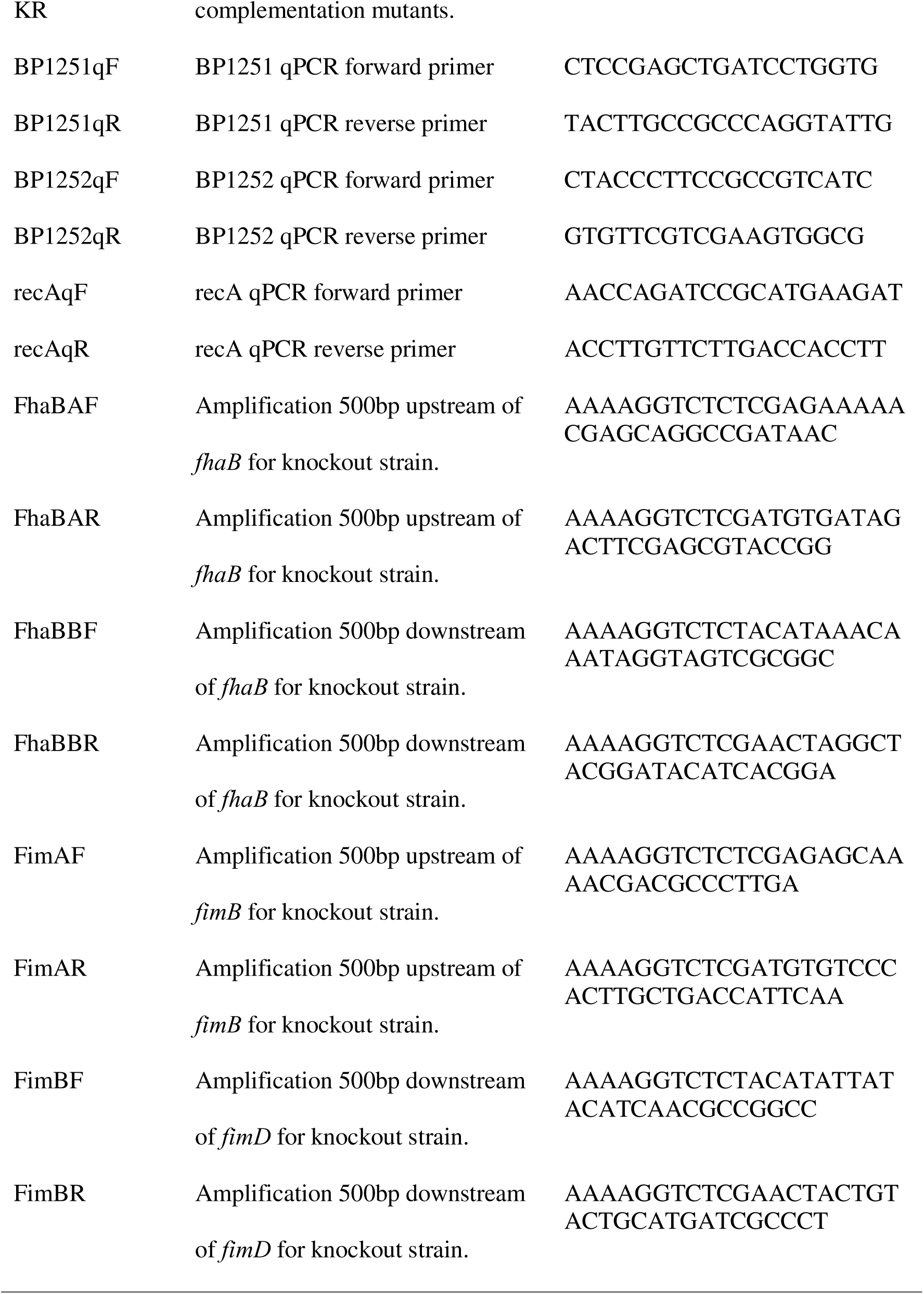
Primers used in this study.

### B. pertussis Conjugations

*E. coli* ST18 was used as the donor to transfer plasmids into *B. pertussis*, using 50μg/mL aminolevulinic acid (ALA) and 5mM MgCl_2_. Streaked plates were cultivated at 37°C for 3-5 days. Knockout mutants were selected using gentamicin (30µg/mL) resistance in the presence of 50mM MgSO_4_. The second homologous recombination event was selected by growth on charcoal agar without MgSO_4_. Gene knockout was confirmed through PCR and nanopore whole-genome sequencing. Complementation of knockout strains was conducted using kanamycin (50µg/ml).

### Growth Curves

*B. pertussis* strains were cultured overnight in SS media at 37°C 200rpm, adjusted to OD_600_ 0.15 and 500μL aliquoted in duplicate into wells of a 24-well microwell plate (Greiner Bio-One). Absorbance at 600nm was measured every hour in a plate reader (Tecan) over three days, with orbital shaking at 37°C.

### Biofilm assay

*B. pertussis* was cultured in SS media overnight, the OD_600_ adjusted to 0.15 and aliquoted in duplicate into wells of a 24-well plate (Greiner Bio-One) and cultured for a further 48 hours at 37°C under static conditions. Media was then carefully removed and each well washed twice with PBS. Wells were then allowed to dry for 15 minutes inside a Class II Biosafety Cabinet. Crystal violet (1% w/v) stain was then applied at 37°C for 30 minutes, followed by two PBS washes, and resuspension of bacterial biofilms in 100% acetic acid. Serial dilutions were performed in PBS and measured at 590nm using a plate reader (Tecan). Experiments were performed in triplicate.

### Tissue Culture

Human respiratory cell lines were cultured for use in infection assays. Two lung/bronchial (A549 and BEAS-2B) and a monocyte (THP-1) cell line were used. THP-1 were differentiated by the addition of 150ng/mL phorbol 12-myristate 13-acetate (PMA) (ThermoFisher Scientific). The epithelial cell lines were cultured according to ATCC guidance for the specific cell line and as adherent cells. All cell lines were maintained until a maximum passage number of 25. A549 and BEAS-2B were cultured in Gibco Dulbecco’s modified eagle’s medium (DMEM) (ThermoFisher Scientific) 10% Gibco Foetal Bovine Serum (FBS, ThermoFisher Scientific), whilst THP-1 were cultured in Gibco RPMI 1640 (ThermoFisher Scientific) 10% FBS.

### Cell Binding Assays

Cell binding assays were performed in 24-well cell culture plates (Greiner Bio-One), with each sample in duplicate, performed across three independent experiments. Assay plates were seeded with the relevant cell line at ∼70% confluency to 5×10^3^ cells per well in 500µL DMEM 10% FBS and then incubated ∼18-24 hours at 37°C, 5% CO_2_. *B. pertussis* colonies were resuspended in FBS-free DMEM, at a multiplicity of infection (MOI) of 1000. Cells were washed twice with FBS-free DMEM and media containing *B. pertussis* strains was added to each well. The *B. pertussis* suspensions were serially diluted and plated on agar to enumerate the inoculations used. The infection was synchronised by centrifugation at 900rpm for 3 minutes and then incubated at 37°C, 5% CO_2_ for 45 minutes. Wells were washed three times with sterile PBS. 500µL MilliQ (Merck) water was added to each well and the adherent cells physically detached. This solution was passed through a 25G needle before 10-fold serial dilution. Each dilution was plated on charcoal agar and incubated for five days before colony enumeration. Colony counts were adjusted for dilution and normalised to WT strain binding. At least three independent experiments were conducted, with duplicate samples in each.

### Cell invasion Assays

24-well plates (Greiner Bio-One) were seeded with 1×10^4^ A549 cells/well in DMEM 10% FBS and incubated for 24 hours. Media was removed and replaced with DMEM (without FBS) containing each bacterial strain. The MOI was adjusted from 1000 to account for differential binding between strains, determined by cell binding assay data. The *B. pertussis* suspensions were serially diluted and plated on agar to enumerate the inoculations used. Plates were incubated at 37°C 5% CO_2_ and samples were harvested at 0h and 24h. At each timepoint, unattached bacteria were removed with the media and wells were washed twice with PBS before media containing gentamicin at 30ug/mL was added to kill remaining extracellular bacteria. After 1h incubation, wells were washed twice with PBS before physical detachment of the immobilised cells and aspiration through a 25G needle. 10-fold serial dilutions were performed and plated on charcoal agar, and colonies counted after five days of incubation at 37°C. Three independent experiments were conducted, with duplicate samples in each.

### Lactate Dehydrogenase Cytotoxicity Assays

Eukaryotic cells at 60-80% confluency were seeded in a 96-well (Greiner Bio-One) at 5×10^3^ cells/well in 100μL 10% FBS Phenol red-free RPMI (ThermoFisher Scientific) and incubated ∼18-24 hours at 37°C 5% CO_2_. *B. pertussis* colonies were resuspended in PBS and adjusted to OD_600_ 1 before diluting to OD_600_ 0.006, representing an MOI of 100. Serial dilutions were performed for MOI 10 and 1. A 5% TritonX-100 positive control was included. Plates were centrifuged at 900rpm for 3 minutes before incubation at 37°C 5% CO_2_ for 3h, 6h, and 24h. 50µL supernatant from each well was aliquoted into 50µL LDH Assay Substrate Buffer (ThermoFisher Scientific) and incubated at 37°C for 30 minutes. The reaction was stopped through addition of the Stop Buffer before measuring absorbance at 490nm and maximum LDH release.

### Mouse Colonisation

C57BL6/j black mice were purchased from Jackson laboratories and maintained in our facilities where they were kept under the care of the employees and veterinarians of Louisiana State University Health - Shreveport Animal Care Facility, Shreveport, LA, (AUP:20-038, AUP:22-031). All experiments were carried out in accordance with all institutional guidelines (AUP:20-0038, AUP:22-031).

For our animal inoculations 4-6 weeks old mice were anesthetized with 5% isoflurane (Attane, # RXISO-250) and when asleep they were intranasally challenged with 50μl of PBS (Gibco, #10010-031) containing 5×10^5^ CFU/mL of Bp536, ΔBP1251, ΔBP1252, and ΔBP1251-1252. Mice were euthanized at different times post-inoculation using 5% CO_2_ followed by cervical dislocation in accordance with the humane endpoints included in our protocols. To perform the organ collection, we used 2ml reinforced homogenizer tubes (VWR, #10158-556), containing 1ml of sterile PBS and a mixture of 0.5 mm and 1.4 mm ceramic beads. Data was collected from at least two independent experiments, with at least four mice per time and condition.

### Bioinformatics

Standalone BLASTp (2.11.0, [26]) was used to for amino acid homology analysis. A manually curated *Bordetella* species protein database was generated from the UniProt database ([27]). The NCBI non-redundant database was also used. BLASTp was conducted with an e-value cut-off of 1×10^5^.

### Statistical Analyses

One-way ANOVA was conducted for statistical analyses across all experiments, followed by pairwise Tukey’s T-tests for between-group testing. P values were represented as <0.05 (*), <0.01 (**), <0.001 (***). Error bars were used to represent Standard Error of the Mean (SEM). Data was compiled and analysed within in Excel and Graphpad Prism 9.0. Graphs were generated in Graphpad Prism 9.0.

## Results

### BP1251 and BP1252 are novel Bordetella spp. putative toxins that are highly conserved amongst Bordetellae

Protein homology was screened using Basic Local Alignment Search Tool for Proteins (BLASTp, [28]) using a manually curated database of *Bordetella* protein sequences, obtained from UniProt [27], and the NCBI non-redundant protein database. The closest annotated homologs of BP1251 were Pertussis toxin (Ptx) subunits 2 (PTx S2, 30% identity and 85% query coverage) and 3 (PTx S3, 32% identity and 83% query coverage), of the PTxB hetero-hexamer. Similar hits were reported when using the non-redundant protein database, with no hits reported outside of the *Bordetella* genus. A single annotated hit was reported for BP1252, with the presence of a PRK15266 domain denoting its closest homolog as Subtilase Cytotoxin subunit B (SubB) of *Salmonella enterica* (25% identity, 90% query coverage). Both proteins, therefore, were homologous to the B components of AB toxins. Both proteins are conserved in the two other classical *Bordetella* species, *B. bronchiseptica* and *B. parapertussis*, with >97% identity and 100% query coverage(Appendix 2), and absent in the other Bordetella species.

### BP1251 and BP1252 do not contribute to biofilm formation in vitro

To explore the general adherence properties of KO mutants, biofilm formation was studied using a crystal violet assay after two days of stationary growth in SS. There were no significant differences in biofilm formation between the mutant strains compared to WT suggesting they do not contribute to in vitro biofilm formation. (Appendix 3).

### BP1251 and BP1252 are required for B. pertussis binding to lung epithelial cells but not for cell invasion or cytotoxicity

To determine the role in adherence to epithelial cells, A549 and BEAS-2B cells were infected with *B. pertussis* mutant strains at an MOI of 1000. Data showed a significant reduction in binding of Δ*bp1251* and Δ*bp1252* strains to A549 cells of 50% (p=0.016) and 60% (p=0.001) (Figure 1a), respectively, relative to WT. Further, complementation of Δ*bp1251* and Δ*bp1252* strains increased binding of *B. pertussis* to A549 cells to above that observed in the WT strain by 57% (p<0.001) and 101% (p<0.001), respectively (Figure 1a). Similarly, both Δ*bp1251* and Δ*bp1252* strains had significantly reduced binding to BEAS-2B bronchial epithelial cells, by 39% (p=0.001) and 36% (p=0.009), respectively (Figure 1b). Complementation of Δ*bp1251* increased BEAS-2B binding, restoring it to levels observed in WT (Figure 1b). Deletion of both *bp1251* and *bp1252*, in the Δ*bp1251-52* strain, also significantly reduced BEAS-2B binding by *B. pertussis* (p<0.001), by 45% (Figure 1b). A dual Δ*bp1251-1252* KO mutant strain also showed reduced binding (Figure 1) to both A549 (35%) and BEAS-2B cells (45%, p=0.0008). Fha and Fim KO mutants were included for comparison and showed 40-60% reduced binding to both A549 (Figure 1a) and BEAS-2B (Figure1b).

**Figure 1.**
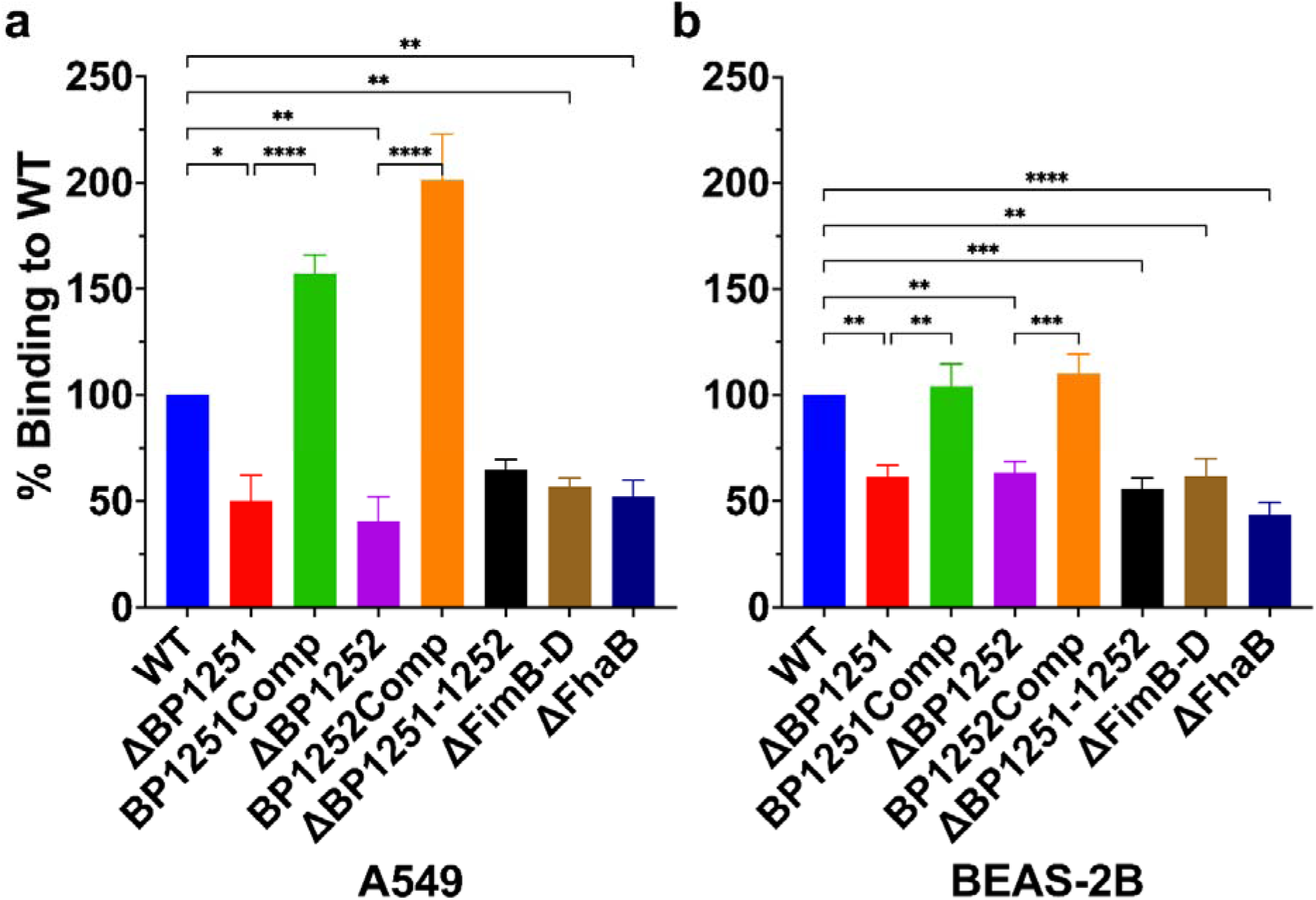
Lung Epithelial Cell Binding of *B. pertussis* Mutant Strains. Binding of BP1251 and BP1252 knockout and complemented strains, and FHA and Fim KO strains, to A549 (**a**) and BEAS-2B (**b**) lung epithelial cells, applied at a multiplicity of infection (MOI) of 1000. Values represent percentage binding to Bp536 (WT) strain. Three independent experiments were performed, with each sample in duplicate. Statistical analyses were performed using one-way ANOVA. Error bars represent SEM.

To determine if BP1251 and BP1252 play a role in invasion of epithelial cells, a cell invasion assay was carried out. The number of bacteria of each strain used was adjusted to compensate for the defect in binding of the mutants and after allowing 45 min for the bacteria to invade, extracellular bacteria were killed by exposure to gentamicin before enumerating intracellular bacteria at 0 and 24h. Across three experiments, initial invasion of A549 cells by *B. pertussis* mutant strains showed no significant differences compared to WT (Figure 2a). There was no significant difference in the number of intracellular bacteria after 24 hours of incubation (Figure 2b), suggesting that there was no effect on proliferation inside the cells.

**Figure 2.**
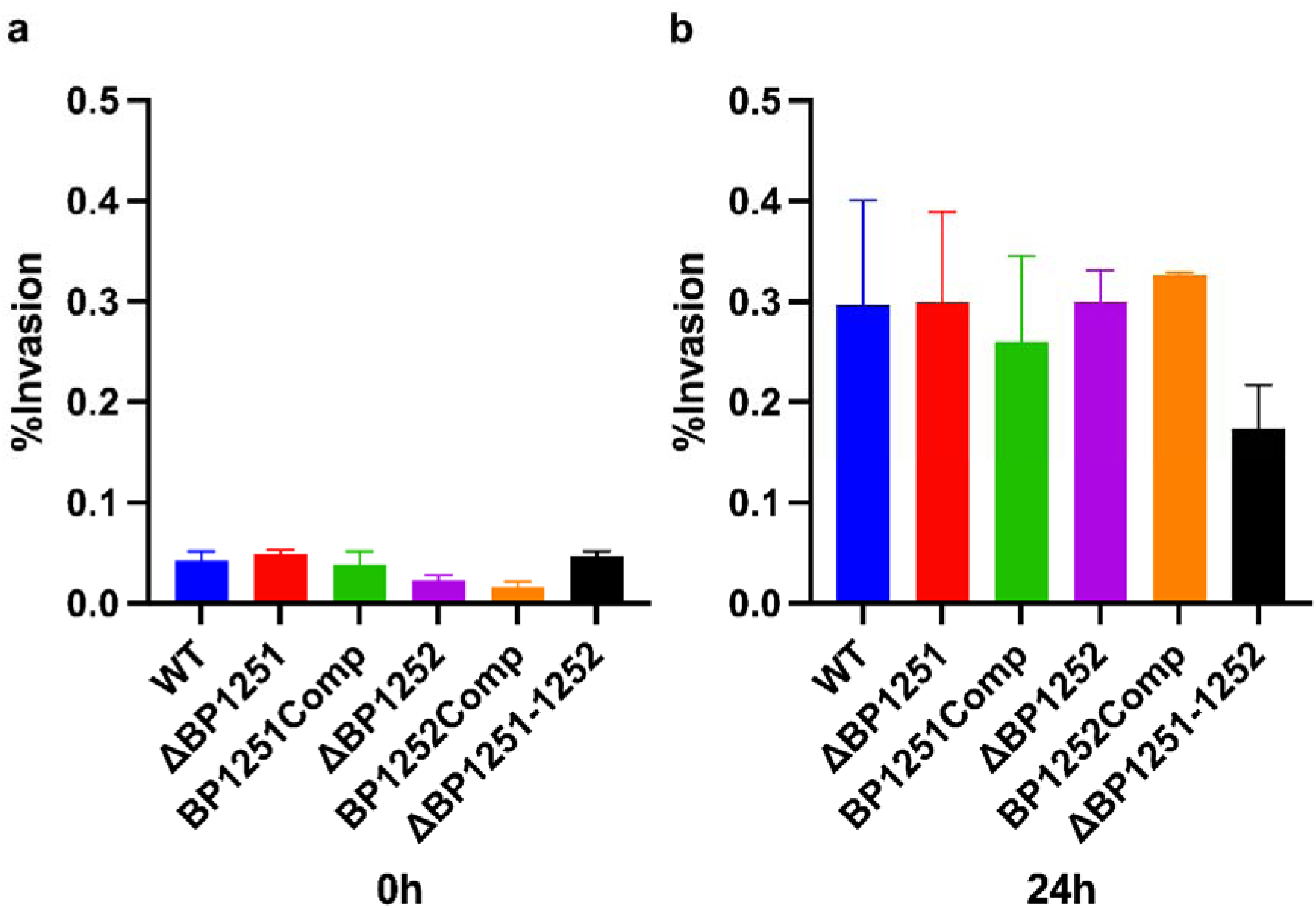
The role of BP1251 and BP1252 in Lung Epithelial Cell Invasion. Invasion of BP1251 and BP1252 *B. pertussis* mutants was investigated at 0h **(a)** and 24h **(b)** timepoints. Values represent percentage of intracellular bacteria compared to inoculum. Statistical analyses were performed using One-way ANOVA. Error bars represent SEM.

Cytotoxicity assays were carried out with the mutant *B. pertussis* strains, using human respiratory tract and macrophage cell lines. Two methods were used, measuring lactate dehydrogenase (LDH) release and propidium iodide fluorescence. When applied to A549 alveolar epithelial cells, all strains demonstrated similar toxicity to WT across three different bacterial concentrations and timepoints (Figure 3), with no significant differences observed. This was observed also in the propidium iodide assay. Cytotoxicity of mutant *B. pertussis* strains to human macrophage cells was also similar to the WT strain (Appendix 4).

**Figure 3.**
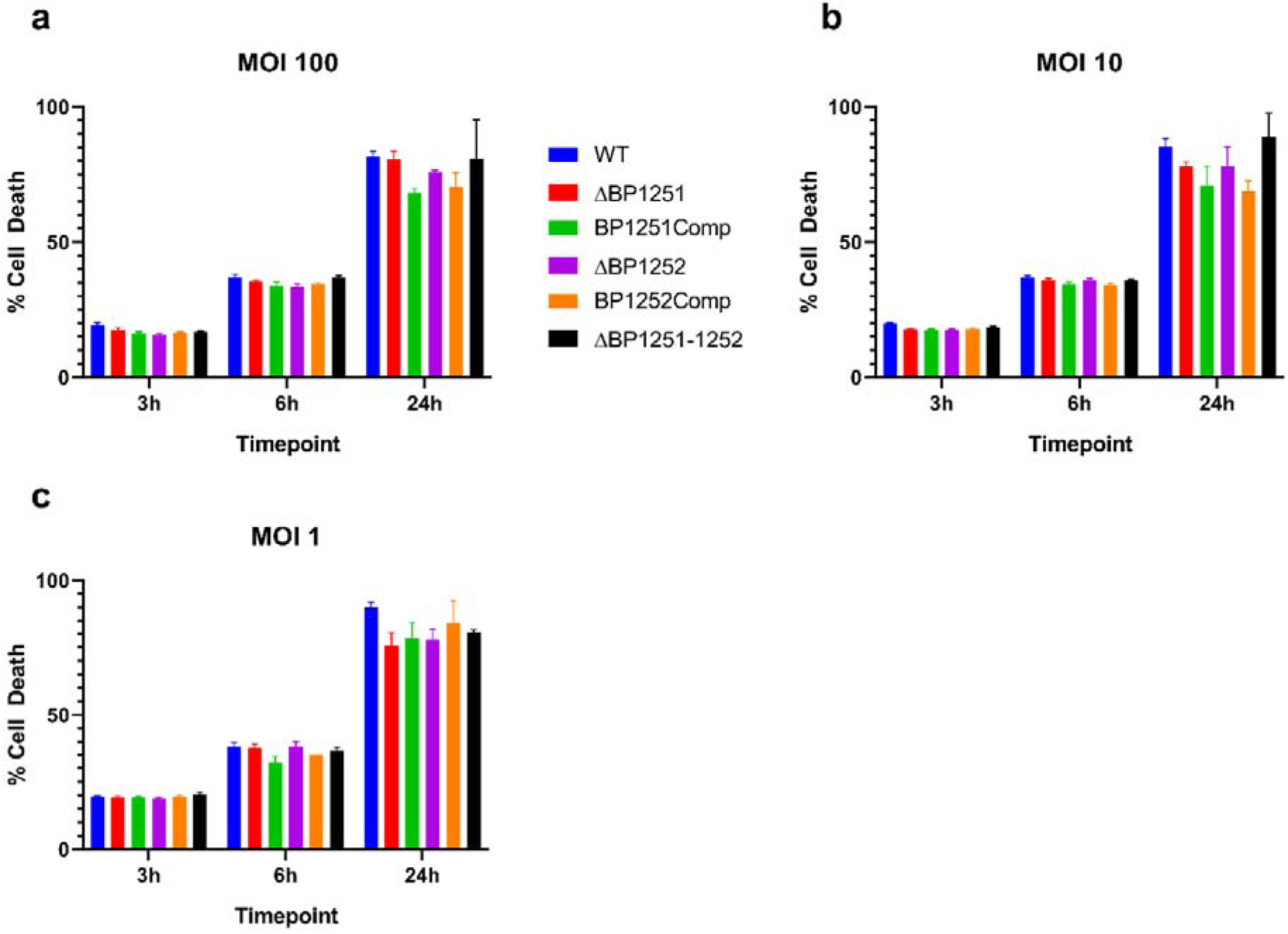
Cytotoxicity of *B. pertussis* Mutant Strains. Cytotoxicity of BP1251 and BP1252 single knockout mutants and their cognate complementation strains to A549 cells, applied at a multiplicity of infection (MOI) of 100 (**a**), 10 (**b**), and 1 (**c**). Lactate dehydrogenase (LDH) was measured at 490nm after 3h, 6h, and 24h incubation. Values represent cell death as a percentage of maximum LDH release. Samples were analysed in triplicate. Statistical analyses were performed using two-way ANOVA. Error bars represent SEM.

### BP1251 and BP1252 promote colonisation of upper respiratory tract

To explore the role of BP1251 and BP1252 *in vivo*, mice were intranasally inoculated with WT and mutant *B. pertussis* strains. Nasal, tracheal, and lung tissue were removed after 0, 0.5, 1, 3, and 7 days-post-infection (dpi), and bacteria enumerated through colony counting. Colonisation of the upper respiratory tract was significantly reduced in the mutant *B. pertussis* strains (Figure 4a). At 1 dpi, significantly fewer Δ*bp1251* (p=0.0003), Δ*bp1252* (p=0.0004), and Δ*bp1251-52* (p=0.0003) were recovered from murine nasal tissue compared to WT (Figure 4b). These reductions in nasal colonisation persisted after 3 dpi (Figure 4c), with significantly lower colonisation by Δ*bp1251* (p=0.0086), Δ*bp1252* (p=0.0095), and Δ*bp1251-1252* (p=0.0091).

**Figure 4.**
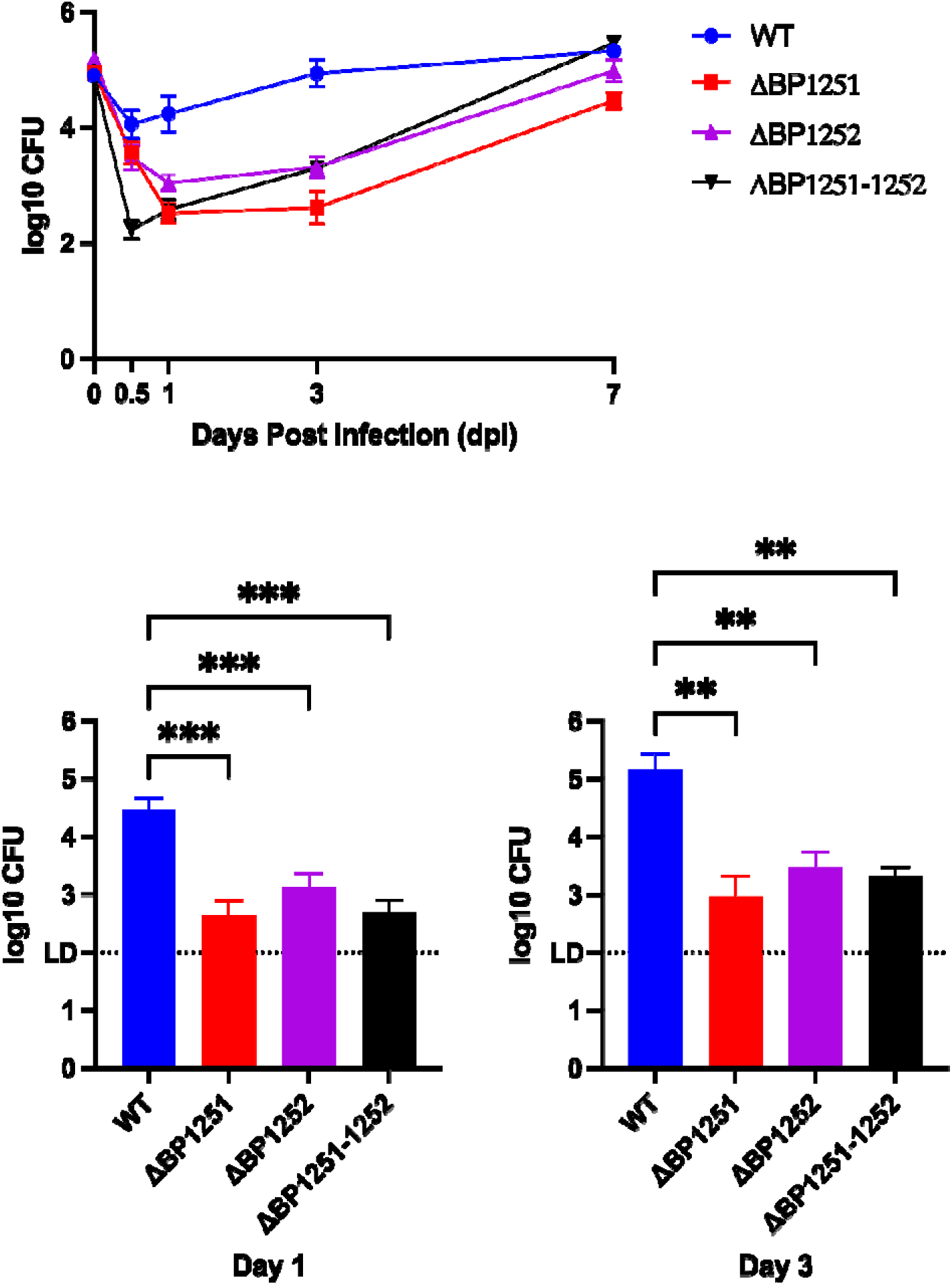
The role of BP1251 and BP1252 in Colonisation of Murine Nasal Cavity. Mutant *B. pertussis* strains were administered to mice nasally and enumerated from nasal cavity tissue across 7 days post-infection **(a).** Bar charts were generated for 1 **(b)** and 3 **(c)** days post-infection. Data is represented as log10 CFU. Statistical analyses were conducted using one-way ANOVA. Error bars represent SEM. Dotted lines mark the limit of detection (LD).

Mutant strains were less impacted in binding to tracheal tissue (Figure 5a) than nasal tissue. Specifically, after 3dpi, no significant reduction in tracheal colonisation was observed in the mutant strains relative to WT (Figure 5c). However, at 1dpi, Δ*bp1252* showed a significant (p=0.03) reduction in colonisation compared to WT (Figure 5c).

**Figure 5.**
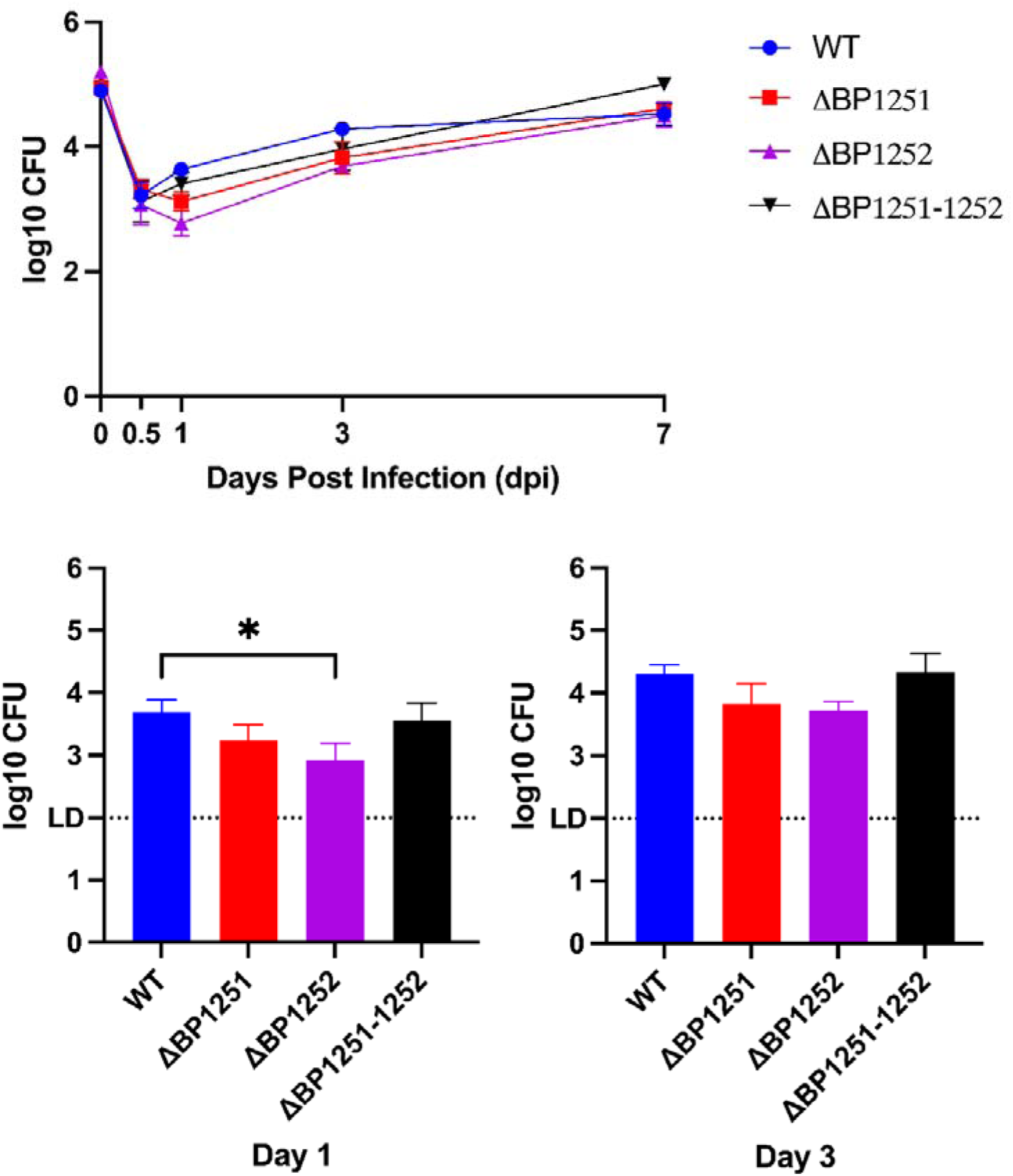
The role of BP1251 and BP1252 in Colonisation of Murine Trachea. Mutant *B. pertussis* strains were administered to mice nasally and enumerated from tracheal tissue across 7 days post-infection **(a).** Bar charts were generated for 1 **(b)** and 3 **(c)** days post-infection. Data is represented as log10 CFU. Statistical analyses were conducted using one-way ANOVA. Error bars represent SEM. Dotted lines mark the limit of detection (LD).

Mutant strains also showed differential lung colonising capabilities (Figure 6a). At 1dpi (Figure 6b) and 3dpi (Figure 6c), all mutant strains showed a significant reduction in colonisation to WT.

**Figure 6.**
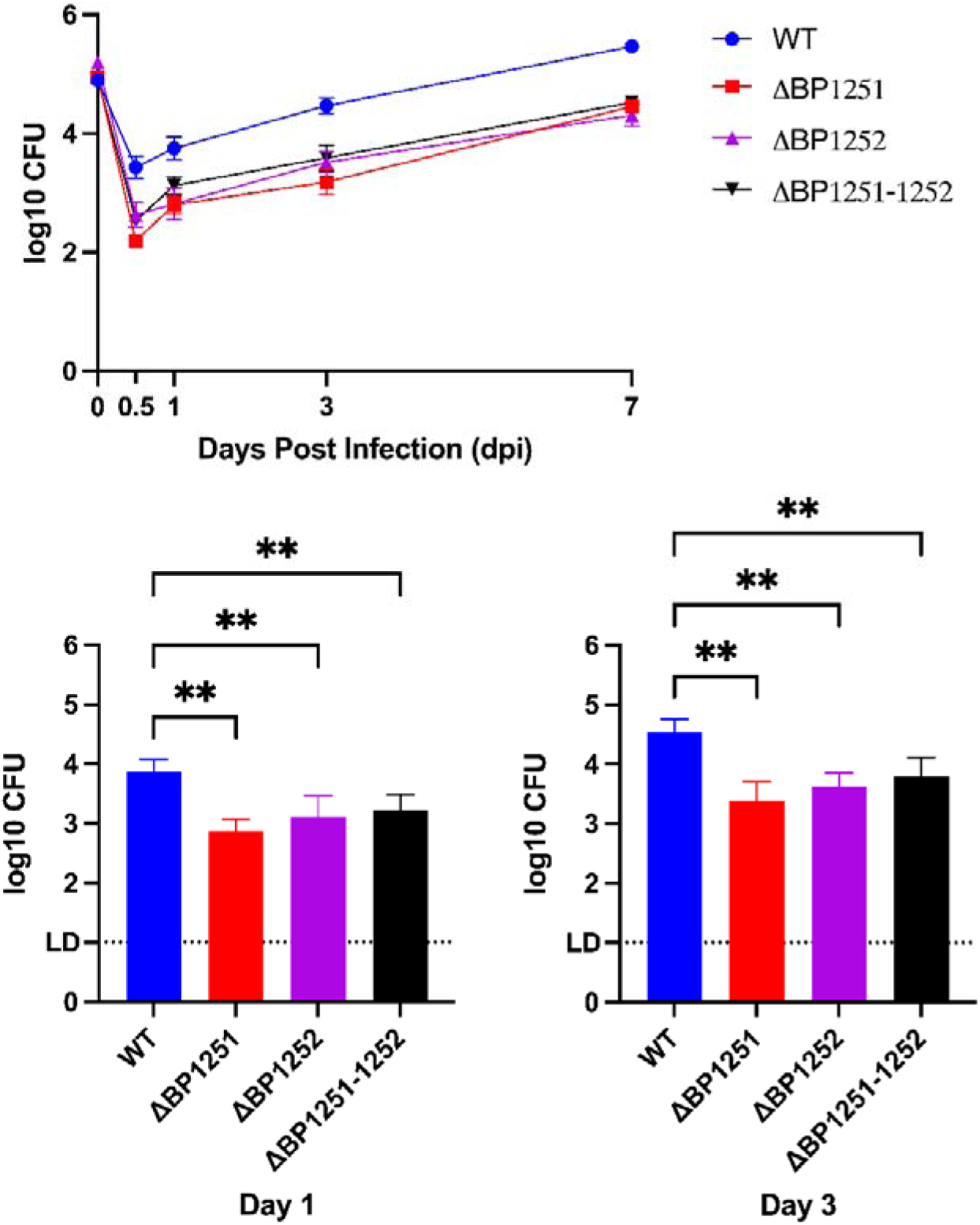
The role of BP1251 and BP1252 in Colonisation of Murine Lungs. Mutant *B. pertussis* strains were administered to mice nasally and enumerated from lung tissue across 7 days post-infection **(a).** Bar charts were generated for 1 **(b)** and 3 **(c)** days post-infection. Data is represented as log10 CFU. Statistical analyses were conducted using one-way ANOVA. Error bars represent SEM. Dotted lines mark the limit of detection (LD).

## Discussion

### BP1251 and BP1252 are Expressed in Virulent and Transmission-specific B. pertussis States

The Bvg regulon is used to characterise different virulence states in *B. pertussis*: avirulent (Bvg-), intermediate (Bvgi), and virulent (Bvg+). The intermediate state is only recently being appreciated as an important part of *B. pertussis* virulence, being attributed to host transmission [19]. Differential gene expression between Bvg states is dictated by the affinity of BvgA binding sites within promoter regions. High affinity binding of the phosphorylated transcriptional activator BvgA facilitates gene expression despite low levels of activated BvgA, such as during the intermediate Bvgi state [29].

RNA sequencing showed that both *bp1251* and *bp1252* were transcribed in Bvg+ and Bvgi conditions, with much lower transcription in Bvg-conditions (data not shown). This suggested that both *bp1251* and *bp1252* are likely to be important for virulence. Their expression within the Bvgi state suggests a potential role in *B. pertussis* transmission, with adhesins such as FHA [29], pertactin [17], and fimbriae [17] being expressed in both Bvg+ and Bvgi states, unlike other, non-adhesin, virulence factors [17,18]. The exception is the expression of Ptx in Bvgi [17], which does not function as a direct adhesin [30,31]. This may be due to the indirect role of PTx in the binding of other adhesins to target cells [31] and thus importance during initial host colonisation.

### BP1251 and BP1252 are Putative Adhesins

The homology of both BP1251 and BP1252 to AB toxin B subunits suggests that these proteins are involved in binding to host cell receptors. Orphan B subunits and their respective homologs have a high coverage between their amino acid sequences but low identity score, due to several regions of conservation (Appendix 5) which may bear functional importance, further supporting the potential binding role of these proteins *in viv* The binding portion, B component, of AB toxins are often complex, hetero-multimeric structures. These portions are responsible for binding to target cell surface receptors and translocating the cognate “A” component toxin to mediate the associated toxicity. For example, PTx is composed of four separate B subunits; S2, S3, S4, and S5 to form a hetero-pentameric B component composed with two S4 subunits [32]. The S1, PTxA, catalytic subunit is an ADP-ribosyltransferase that mediates toxicity to target cells via uncoupling of G-protein signalling [33]. Other examples of AB5 toxins include cholera toxin [34], Shiga toxin [35] and subtilase cytotoxin of Shiga-toxigenic *E. coli* [36]. Given the genomic co-localisation of *bp1251* and *bp1252*, their similar Bvg-regulation profiles, and their assignment as orphan B subunits, it can be hypothesised that these proteins may form a multimeric complex. Furthermore, the fact that both, *bp1251* and *bp1252*, have homologs in the classical *Bordetella* species; *B. parapertussis* and *B. bronchiseptica* (Appendix 2) indicates that these proteins were conserve during *Bordetella* spp. speciation process suggesting an important role for BP1251 and BP1252.

It is noteworthy that another co-localised gene, *bp1253*, has been previously studied by Moramarco et al. [37], who ascertained its role as a LONELY GUY (LOG) enzyme possessing phosphoribohydrolase activity. Preliminary RNAseq data suggested that *bp1253* transcription is not bvg-regulated (data not shown), with equivalent transcription across the three Bvg states. It is also therefore unlikely to be co-transcribed with the Bvg-regulated genes. Further, an initial experiment found no impact on cytotoxicity or adhesion in a BP1253 KO strain (data not shown). Given these data, we chose to continue our experiments only with *bp1251* and *bp1252*.

### Deletion of bp1251 and bp1252 Reduces Lung Epithelial Cell Binding

Mostly, B components, including both subtilase cytotoxin [38,39] and pertussis toxin [40,41], possess binding specificity to sialylated glycans/glycoproteins on cell surfaces. The binding of these B components to their targets facilitates endocytosis of the entire AB complex into the target cell and consequent signalling dysregulation via the A toxin [42]. Several studies have highlighted the retained receptor binding function of B subunits independent of the A subunit activity [21,22], including PTx S2 and S3 subunits [43,44]. Given the apparent absence of a cognate “A” toxin but sequence similarities to B subunits, it was proposed that both BP1251 and BP1252 may facilitate a binding interaction similar to these homologs. Indeed, our results showed a significant reduction in binding to both lung cells of both Δ*bp1251* and Δ*bp1252* mutant strains compared to WT, with a further increased binding when overexpressing the relative proteins in complemented strains. Brief examination of conserved regions between BP1251 and pertussis toxin S2/S3 (Appendix 5) found homology in confirmed binding sites, including L20-T21[43,45] and R35 [43,45–47]. Interestingly these sites dictate the specificity of binding to the sialated glycan/glycoprotein, which might contribute to host cell specific phenotypes. In fact, our results show some differences in the binding affinity of these proteins to bronchial and alveolar epithelial cells, which could be contributing to different steps during the infectious process. Such tissue-dependent colonisation has been observed previously in *B. pertussis* adhesins [30], and in other respiratory pathogens [48], and is likely the result of slight differences in epithelial cell surface compositions as is described in other infection models [48]

Interestingly, a dual KO mutant showed no cumulative defect in binding (Figure 1). This could suggest that BP1251 and BP1252 are co-dependent, with loss of either protein resulting in the same phenotype, which will further support our hypothesis that both proteins form a multimeric complex that might allow for infection of specific tissues. It is worth noting that complementation was done with a low copy number plasmid, which likely results in increased amounts of BP1251 and BP1252 produced, our results showed a significant increase in binding to A549 cells, providing further evidence that these proteins function as very effective adhesins, even in low copy number.

### BP1251 and BP1252 do not Facilitate Epithelial Cell Invasion

Invasion of host epithelial cells by *B. pertussis in situ* may facilitate immune evasion, characteristic of chronic Pertussis infections, and appears to enable more established disease [49]. Neither BP1251 nor BP1252 appear to mediate bacterial invasion of epithelial cells (Figure 2). No significant differences were observed between WT and mutant strains after 1h incubation (Figure 2a).

An expected increase in intracellular bacteria was observed after 24h incubation in all strains although there was no significant difference to WT (Figure 2b), suggesting that there is no function in intracellular proliferation of these proteins. It should be noted that adhesion and invasion properties are not necessarily concomitant, with FHA [50,51] but not pertactin [51,52] facilitating epithelial cell entry.

### BP1251 and BP1252 are not cytotoxic

Cytotoxicity properties of BP1251 and BP1252 were explored due to the intrinsic toxin translocation properties of AB toxin B components. However, deletion of *bp1251* or *bp1252* yielded no significant differences in either lung epithelial (Figure 3) or macrophage cytotoxicity (Appendix 4). These findings were not surprising, as no A component of a toxin was identified in our bioinformatic analysis, leading us to conclude that neither BP1251 nor BP1252 facilitate toxin translocation and thus may suggest that neither belong to novel AB toxins, but rather may be classified as orphan B subunits.

### BP1251 and BP1252 are Important for Colonisation of the Respiratory Tract

Based on our previous data indicating a role for BP1251 and BP1252 in adhesion to alveolar lung cells, we decided to further explore the role of these proteins *in vivo* utilizing the murine model. We intranasally inoculated mice with WT and mutant *B. pertussis* strains to best mimic the natural infection route, and key respiratory tract areas examined at 0, 0.5, 1, 3, and 7dpi timepoints. In the upper respiratory tract, closest to the inoculation site, both Δ*bp1251,* Δ*bp1252*, and the double KO mutant strain showed significantly reduced binding compared to WT (Figure 4). Such large reductions in nasal tissue colonisation (Figure 4c) reflect the observed binding reductions of mutant strains during *in vitro* adhesion assays (Figure 1). These results are highly relevant as this can suggest that these proteins play critical roles during nasal colonisation, which is one of the major issues that we are confronting due to the lack of nasal protection with the current vaccine.

Despite significant reductions of Δ*bp1252* also in tracheal tissues at 1dpi (Figure 5b), similar reductions were not observed in the Δ*bp1251* mutant. This reflects both the differential, tissue-specific, binding of adhesins, and the potential differences between BP1251 and BP1252.

Colonisation of lung tissue was more reflective of the data observed in nasal tissues, with a more significantly reduced binding of mutant strains (Figure 6), indicating that these putative adhesins contribute to lung colonisation. This further support our *in vitro* data revealing a critical role for BP1251 and BP1252 in binding to host cells.

Given the data in this study, we propose that BP1251 and BP1252 be re-named OtbA (BP1251) and OtbB (BP1252) for Orphan Toxin B subunit.

## Conclusion

The identification and characterisation of *B. pertussis* virulence factors is paramount to improving our understanding of its disease, and thus allowing improvements to current vaccination approaches. Many research efforts focus on developing vaccines that prevent nasal colonization subsequently halting transmission. Our two herein characterized proteins, BP1251 and BP1252, are strongly involved in the establishment colonisation of murine nasal tissue, particularly BP1252, and in colonisation of lung tissue, suggesting that they could be further investigated for the addition of these proteins in the current aP vaccine with the goal of decrease the current transmission rates which are rapidly increasing. Interestingly, these proteins appear to be critical for the initial establishment of infection but they do not appear to play a role in the subsequent infection as demonstrated by the invasion assay and mouse infections. Indeed, it is this transmissive state and thus upper respiratory tract colonisation which is often attributed to the increasing prevalence of Pertussis [9,10]. Specifically, enhanced transmission associated with increased prevalence, but no increase in disease severity [53], reflects the proposed inability of aP vaccines to prevent upper respiratory tract colonisation [9,10,53]. It is therefore important to identify virulence factors, such as BP1251 and BP1252, that may facilitate upper respiratory tract colonisation in order to understand host colonisation, and thus attenuate disease transmission and potentiation.

Our data suggests that BP1251/1252 adhesin we have identified is as important as FHA and Fimbria in colonisation of epithelial cells by *B. pertussis*, with a lack of any one of these adhesins resulting in a similar phenotype. The colonisation of respiratory epithelial cells by B. *pertussis* is a result of the action of these three adhesins and that each are necessary but not sufficient to facilitate binding. The binding to the epithelial cells appears to be the main function of these adhesins as they do not appear to have a general adherence function such as biofilm formation in abiotic surfaces. Still further investigations will need to confirm that they do not contribute to biofilm in vivo, however, our data points towards a more specific role for these adhesins in cell specific binding.

## Acknowledgements

We would like to acknowledge the support of the funding bodies which had no role in the study design, data collection, data analysis, or data interpretation.

## Appendix

**Appendix 1.**
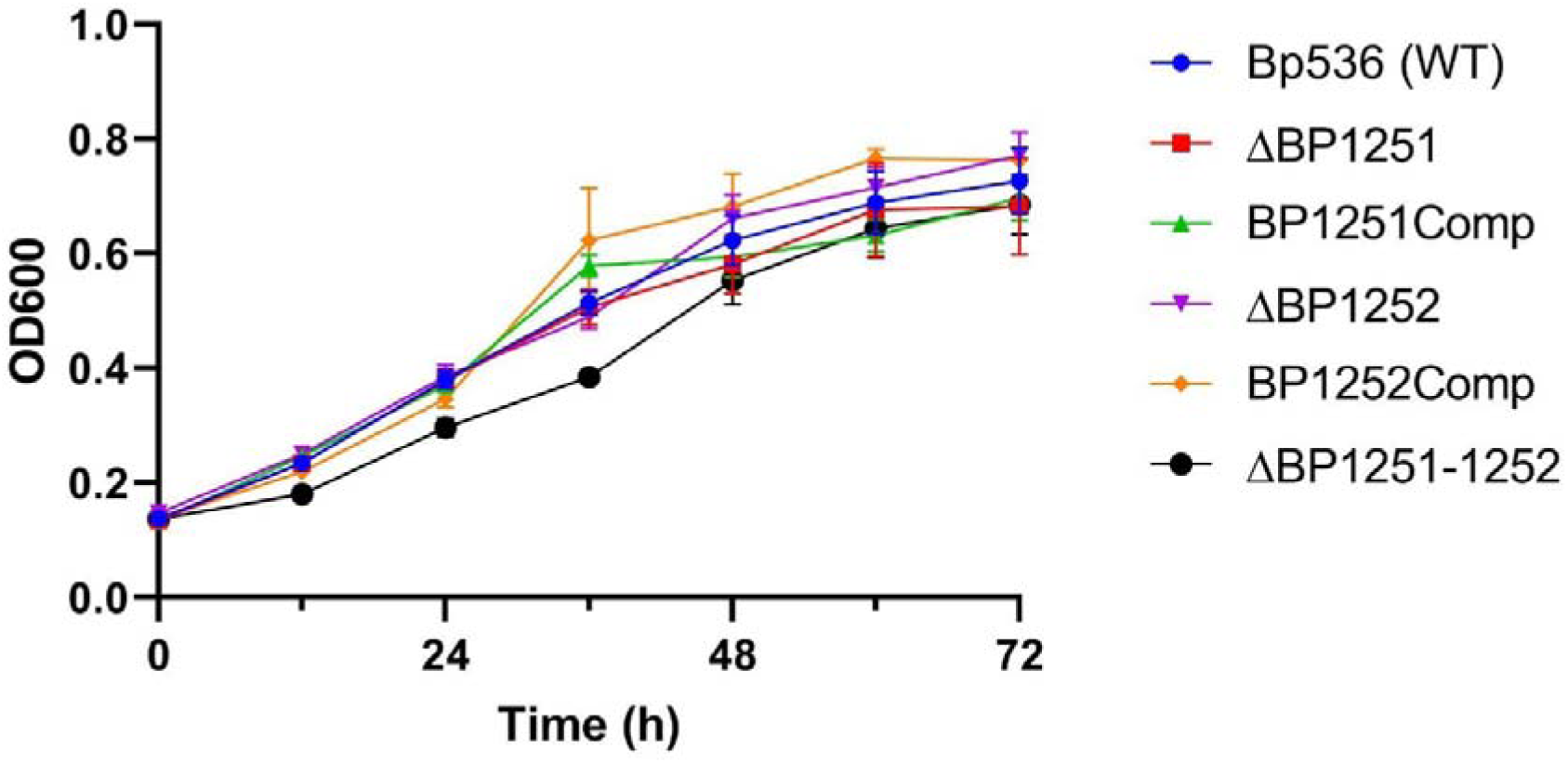
Mutant Strain Growth Curves. Growth of *B. pertussis* strains in Stainer-Scholte broth at 37°C. Error bars represent standard error.

**Appendix 2.**
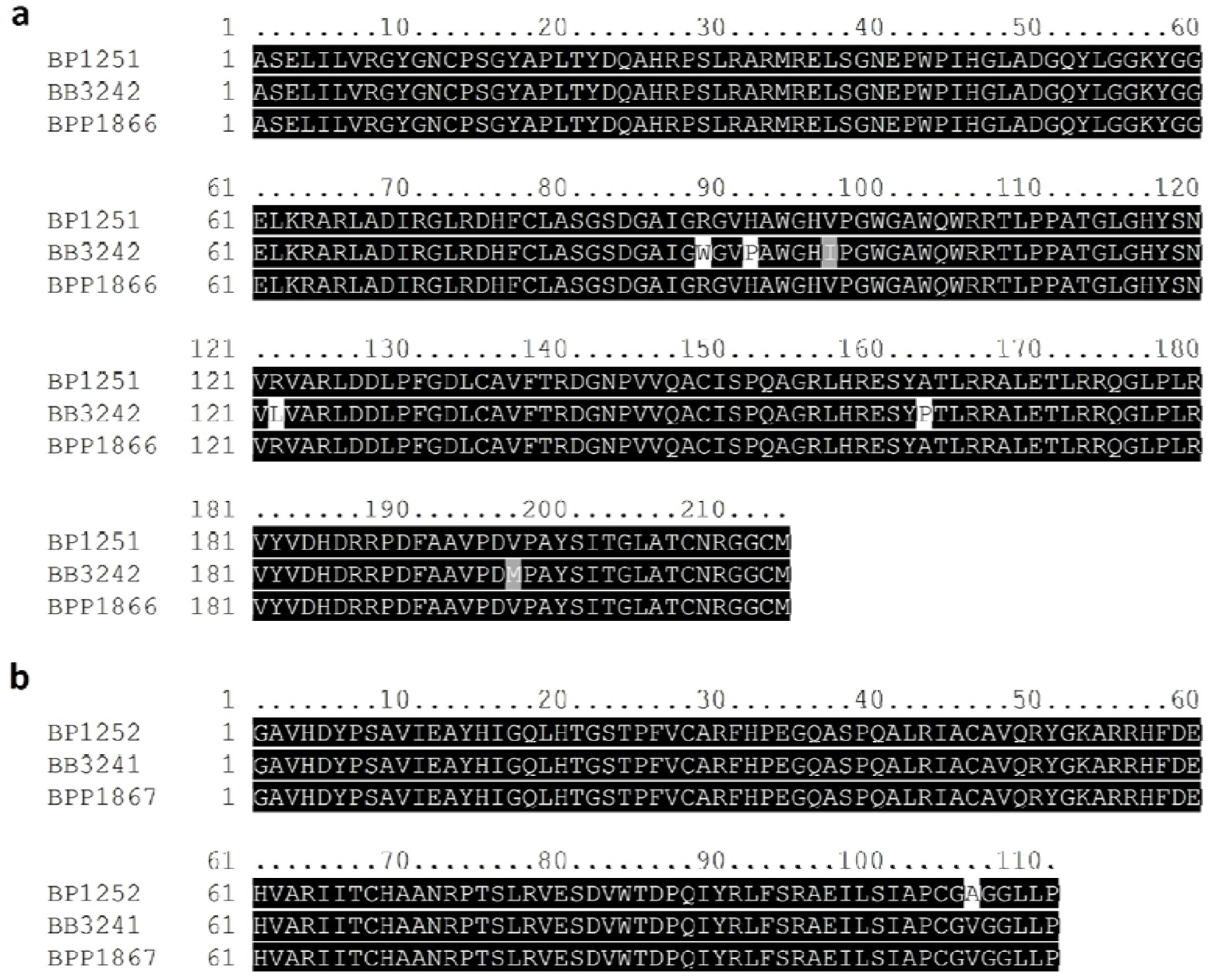
Conservation of BP1251 and BP1252 Within the Classical *Bordetella*. Alignment of BP1251 **(a)** and BP1252 **(b)** with their respective homologs in *B. bronchiseptica* (BB) and *B. parapertussis* (BPP).

**Appendix 3.**
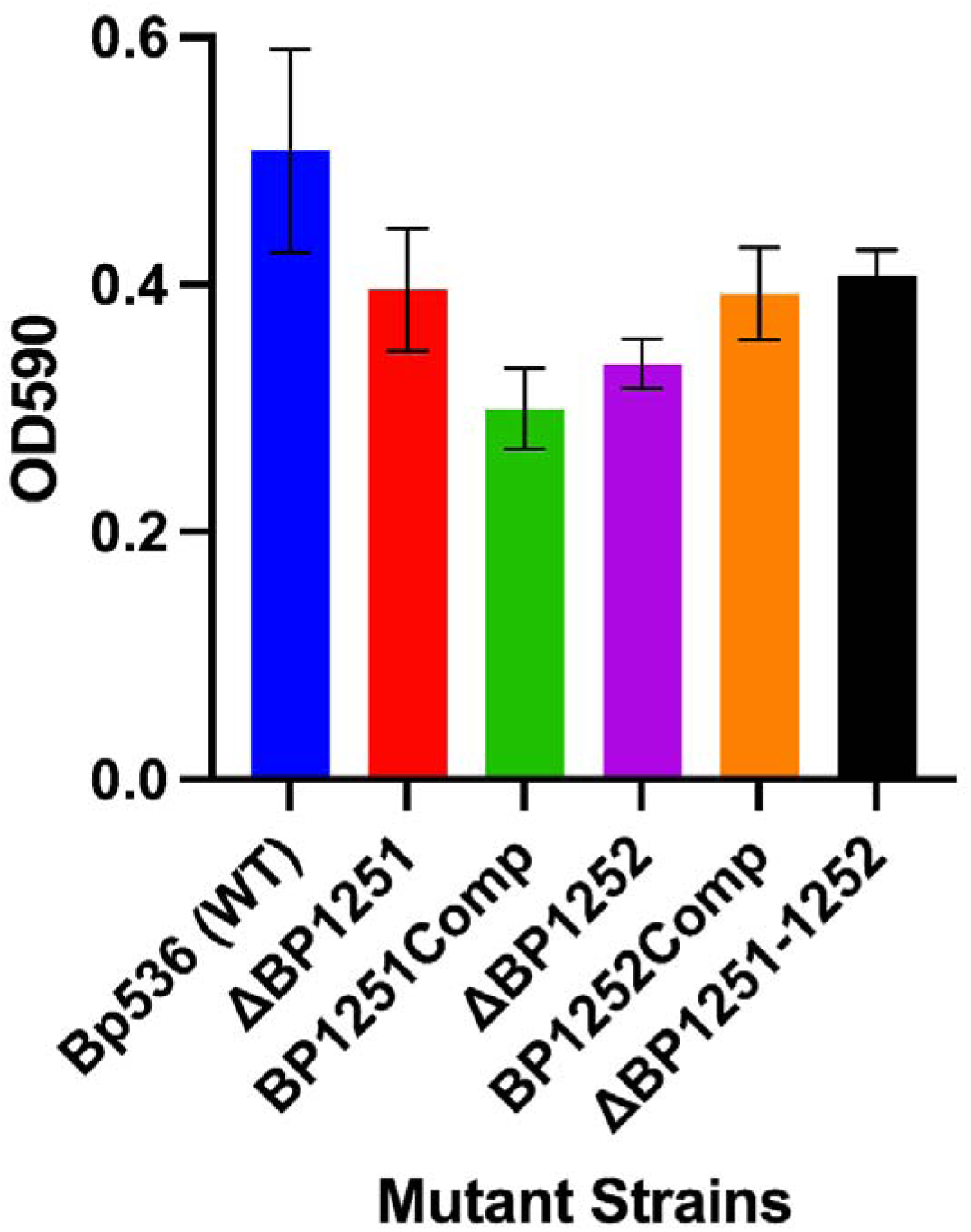
*B. pertussis* Mutant Biofilm Assay. *B. pertussis* strains were cultured and stained using Crystal Violet (1%). Absorbance was recorded at 590nm. No significant difference observed using one-way ANOVA.

**Appendix 4.**
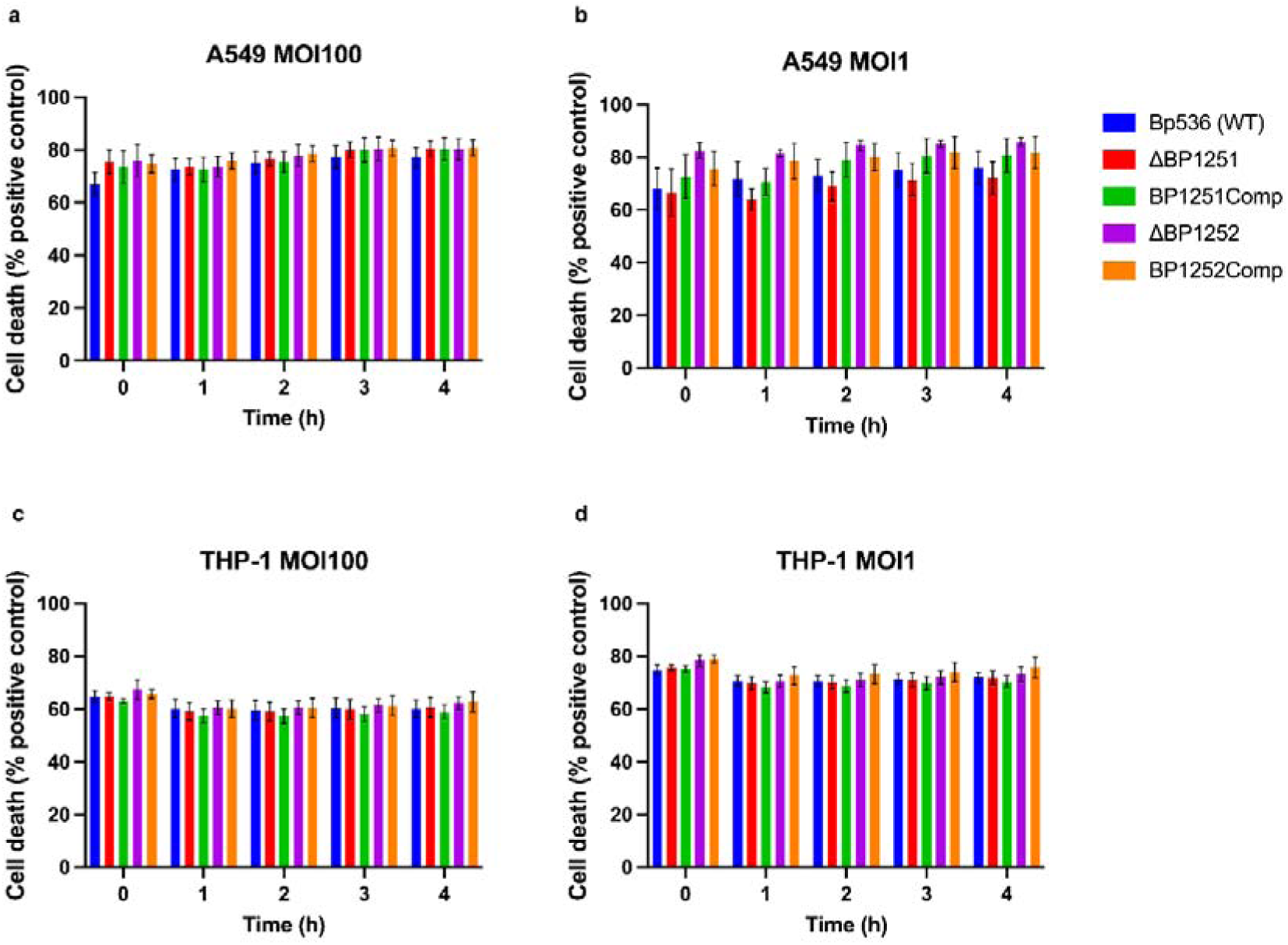
Propidium Iodide Cytotoxicity Assay. Propidium iodide was used to measure cytotoxicity of mutant strains to A549 and THP-1 cells. Fluorescence was measured at 535nm excitation and 624nm emission. Values were adjusted to percentage of WT.

**Appendix 5.**
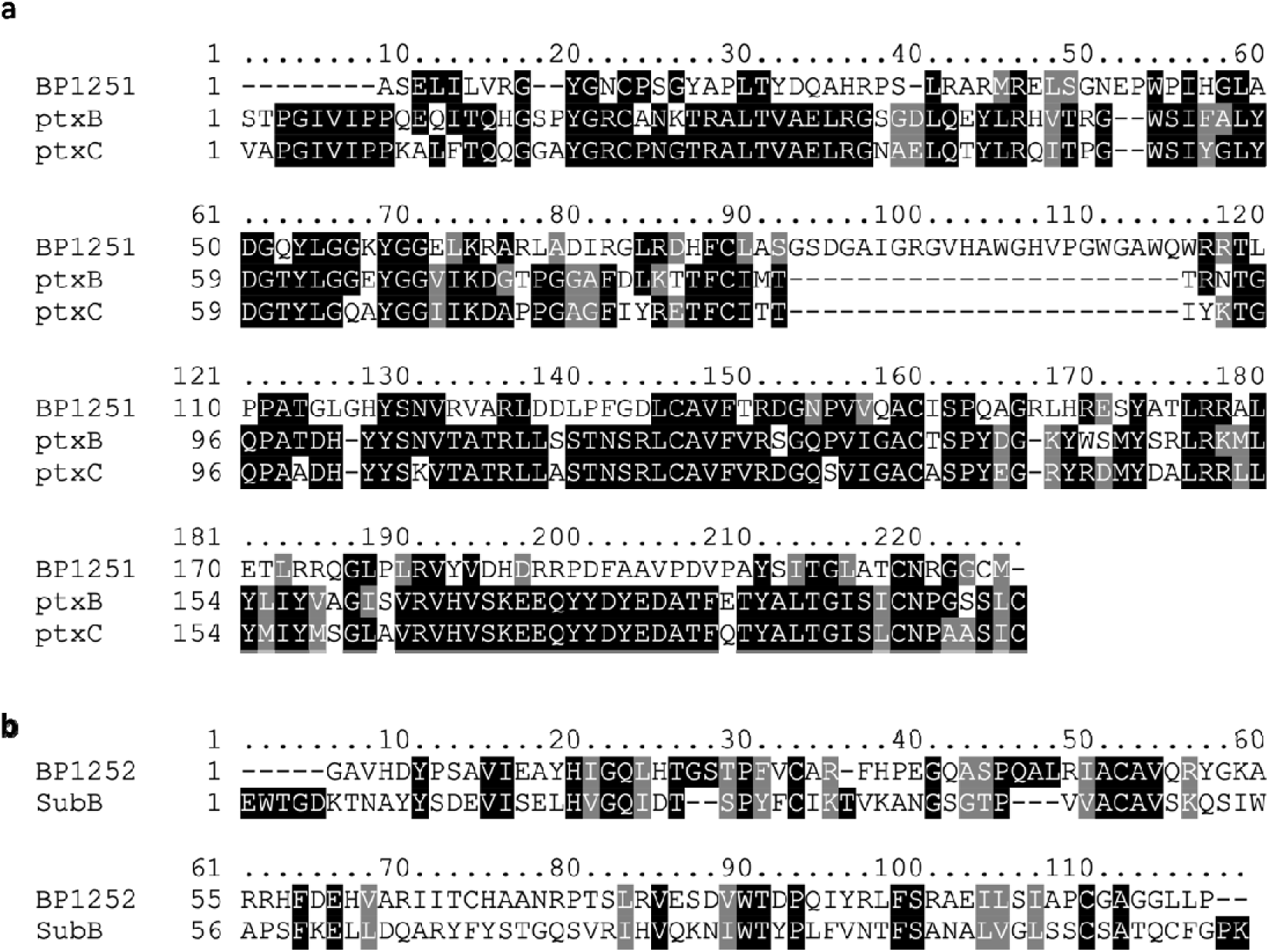
Multiple Sequence Alignment of BP1251 and BP1252 and their Homologs. Mature amino acid sequences were aligned with MEGAX, using Muscle algorithm with default parameters. Boxshade was used to generate the figure, and the conserved regions (CR) annotated manually.

## References

[1] Donnelly S, Loscher CE, Lynch MA, et al. Whole-cell but not acellular pertussis vaccines induce convulsive activity in mice: evidence of a role for toxin-induced interleukin-1beta in a new murine model for analysis of neuronal side effects of vaccination. Infect Immun. 2001;69:4217–4223.

[2] Armstrong ME, Loscher CE, Lynch MA, et al. IL-1β-dependent neurological effects of the whole cell pertussis vaccine: a role for IL-1-associated signalling components in vaccine reactogenicity. J Neuroimmunol. 2003;136:25–33.

[3] Olin P, Gustafsson L, Barreto L, et al. Declining pertussis incidence in Sweden following the introduction of acellular pertussis vaccine. Vaccine. 2003;21:2015–2021.

[4] Vickers D, Ross AG, Mainar-Jaime RC, et al. Whole-cell and acellular pertussis vaccination programs and rates of pertussis among infants and young children. CMAJ. 2006;175:1213–1217.

[5] Soumana IH, Linz B, Dewan KK, et al. Modeling Immune Evasion and Vaccine Limitations by Targeted Nasopharyngeal Bordetella pertussis Inoculation in Mice. Emerg Infect Dis. 2021;27:2107–2116.

[6] Wilk MM, Borkner L, Misiak A, et al. Immunization with whole cell but not acellular pertussis vaccines primes CD4 TRM cells that sustain protective immunity against nasal colonization with Bordetella pertussis. Emerg Microbes Infect. 2019;8:169–185.

[7] Bouchez V, Guillot S, Landier A, et al. Evolution of Bordetella pertussis over a 23-year period in France, 1996 to 2018. Euro Surveill. 2021;26.

[8] Lesne E, Cavell BE, Freire-Martin I, et al. Acellular Pertussis Vaccines Induce Anti-pertactin Bactericidal Antibodies Which Drives the Emergence of Pertactin-Negative Strains. Front Microbiol. 2020;11:2108.

[9] Dubois V, Chatagnon J, Thiriard A, et al. Suppression of mucosal Th17 memory responses by acellular pertussis vaccines enhances nasal Bordetella pertussis carriage. NPJ Vaccines. 2021;6:6.

[10] Holubová J, Staněk O, Brázdilová L, et al. Acellular Pertussis Vaccine Inhibits Bordetella pertussis Clearance from the Nasal Mucosa of Mice. Vaccines (Basel). 2020;8.

[11] Ring N, Abrahams JS, Bagby S, et al. How Genomics Is Changing What We Know About the Evolution and Genome of Bordetella pertussis. Adv Exp Med Biol. 2019;1183:1–17.

[12] Carriquiriborde F, Regidor V, Aispuro PM, et al. Rare Detection of Bordetella pertussis Pertactin-Deficient Strains in Argentina. Emerg Infect Dis. 2019;25:2048–2054.

[13] Esposito S, Stefanelli P, Fry NK, et al. Pertussis Prevention: Reasons for Resurgence, and Differences in the Current Acellular Pertussis Vaccines. Front Immunol. 2019;10:1344.

[14] Ring N, Abrahams JS, Bagby S, et al. How Genomics Is Changing What We Know About the Evolution and Genome of *Bordetella pertussis*. In: Fedele G, Ausiello CM, editors. PERTUSSIS INFECTION AND VACCINES: ADVANCES IN MICROBIOLOGY, INFECTIOUS DISEASES AND PUBLIC HEALTH, VOL 12. 2019. p. 1–17.

[15] Carriquiriborde F, Regidor V, Aispuro PM, et al. Rare Detection of *Bordetella pertussis* Pertactin-Deficient Strains in Argentina. Emerg Infect Dis. 2019;25:2048–2054.

[16] Esposito S, Stefanelli P, Fry NK, et al. Pertussis Prevention: Reasons for Resurgence, and Differences in the Current Acellular Pertussis Vaccines. Front Immunol. 2019;10.

[17] van Beek LF, de Gouw D, Eleveld MJ, et al. Adaptation of Bordetella pertussis to the Respiratory Tract. J Infect Dis. 2018;217:1987–1996.

[18] de Gouw D, Hermans PWM, Bootsma HJ, et al. Differentially expressed genes in Bordetella pertussis strains belonging to a lineage which recently spread globally. PLoS One. 2014;9:e84523.

[19] Vergara-Irigaray N, Chávarri-Martínez A, Rodríguez-Cuesta J, et al. Evaluation of the role of the Bvg intermediate phase in Bordetella pertussis during experimental respiratory infection. Infect Immun. 2005;73:748–760.

[20] Locht C, Antoine R, Raze D, et al. Bordetella pertussis from functional genomics to intranasal vaccination. Int J Med Microbiol. 2004;293:583–588.

[21] Aman AT, Fraser S, Merritt EA, et al. A mutant cholera toxin B subunit that binds GM1-ganglioside but lacks immunomodulatory or toxic activity. Proc Natl Acad Sci U S A. 2001;98:8536–8541.

[22] Richards CM, Aman AT, Hirst TR, et al. Protective mucosal immunity to ocular herpes simplex virus type 1 infection in mice by using Escherichia coli heat-labile enterotoxin B subunit as an adjuvant. J Virol. 2001;75:1664–1671.

[23] Hill BC, Baker CN, Tenover FC. A simplified method for testing Bordetella pertussis for resistance to erythromycin and other antimicrobial agents. J Clin Microbiol. 2000;38:1151–1155.

[24] Stainer DW, Scholte MJ. A simple chemically defined medium for the production of phase I Bordetella pertussis. J Gen Microbiol. 1970;63:211–220.

[25] Thoma S, Schobert M. An improved Escherichia coli donor strain for diparental mating. FEMS Microbiol Lett. 2009;294:127–132.

[26] Camacho C, Coulouris G, Avagyan V, et al. BLAST+: architecture and applications. BMC Bioinformatics. 2009;10:421.

[27] UniProt Consortium. UniProt: the Universal Protein Knowledgebase in 2023. Nucleic Acids Res. 2023;51:D523–D531.

[28] Altschul SF, Gish W, Miller W, et al. Basic local alignment search tool. J Mol Biol. 1990;215:403–410.

[29] Deora R, Bootsma HJ, Miller JF, et al. Diversity in the Bordetella virulence regulon: transcriptional control of a Bvg-intermediate phase gene. Mol Microbiol. 2001;40:669–683.

[30] van den Berg BM, Beekhuizen H, Willems RJ, et al. Role of Bordetella pertussis virulence factors in adherence to epithelial cell lines derived from the human respiratory tract. Infect Immun. 1999;67:1056–1062.

[31] Alonso S, Pethe K, Mielcarek N, et al. Role of ADP-ribosyltransferase activity of pertussis toxin in toxin-adhesin redundancy with filamentous hemagglutinin during Bordetella pertussis infection. Infect Immun. 2001;69:6038–6043.

[32] Stein PE, Boodhoo A, Armstrong GD, et al. Structure of a pertussis toxin-sugar complex as a model for receptor binding. Nat Struct Biol. 1994;1:591–596.

[33] Plaut RD, Scanlon KM, Taylor M, et al. Intracellular disassembly and activity of pertussis toxin require interaction with ATP. Pathog Dis. 2016;74.

[34] Gill DM, Meren R. ADP-ribosylation of membrane proteins catalyzed by cholera toxin: basis of the activation of adenylate cyclase. Proc Natl Acad Sci U S A. 1978;75:3050–3054.

[35] Utskarpen A, Slagsvold HH, Dyve AB, et al. SNX1 and SNX2 mediate retrograde transport of Shiga toxin. Biochem Biophys Res Commun. 2007;358:566–570.

[36] Paton AW, Beddoe T, Thorpe CM, et al. AB5 subtilase cytotoxin inactivates the endoplasmic reticulum chaperone BiP. Nature. 2006;443:548–552.

[37] Moramarco F, Pezzicoli A, Salvini L, et al. A LONELY GUY protein of Bordetella pertussis with unique features is related to oxidative stress. Sci Rep. 2019;9:17016.

[38] Khan N, Sasmal A, Khedri Z, et al. Sialoglycan-binding patterns of bacterial AB5 toxin B subunits correlate with host range and toxicity, indicating evolution independent of A subunits. J Biol Chem. 2022;298:101900.

[39] Day CJ, Paton AW, Higgins MA, et al. Structure aided design of a Neu5Gc specific lectin. Sci Rep. 2017;7:1495.

[40] Heerze LD, Smith RH, Wang N, et al. Utilization of sialic acid-binding synthetic peptide sequences derived from pertussis toxin as novel anti-inflammatory agents. Glycobiology. 1995;5:427–433.

[41] Millen SH, Watanabe M, Komatsu E, et al. Single Amino Acid Polymorphisms of Pertussis Toxin Subunit S2 (PtxB) Affect Protein Function. PLoS One. 2015;10:e0137379.

[42] el Bayâ A, Linnemann R, von Olleschik-Elbheim L, et al. Endocytosis and retrograde transport of pertussis toxin to the Golgi complex as a prerequisite for cellular intoxication. Eur J Cell Biol. 1997;73:40–48.

[43] Saukkonen K, Burnette WN, Mar VL, et al. Pertussis toxin has eukaryotic-like carbohydrate recognition domains. Proc Natl Acad Sci U S A. 1992;89:118–122.

[44] Rossjohn J, Buckley JT, Hazes B, et al. Aerolysin and pertussis toxin share a common receptor-binding domain. EMBO J. 1997;16:3426–3434.

[45] Rozdzinski E, Burnette WN, Jones T, et al. Prokaryotic peptides that block leukocyte adherence to selectins. J Exp Med. 1993;178:917–924.

[46] Tallett A, Seabrook RN, Irons LI, et al. Localisation of a receptor-recognition domain on the S3 subunit of pertussis toxin by peptide mapping. Eur J Biochem. 1993;211:743–748.

[47] van’t Wout J, Burnette WN, Mar VL, et al. Role of carbohydrate recognition domains of pertussis toxin in adherence of Bordetella pertussis to human macrophages. Infect Immun. 1992;60:3303–3308.

[48] Hillyer P, Shepard R, Uehling M, et al. Differential Responses by Human Respiratory Epithelial Cell Lines to Respiratory Syncytial Virus Reflect Distinct Patterns of Infection Control. J Virol. 2018;92.

[49] Baroli CM, Gorgojo JP, Blancá BM, et al. Bordetella pertussis targets the basolateral membrane of polarized respiratory epithelial cells, gets internalized, and survives in intracellular locations. Pathog Dis. 2023;81.

[50] Ishibashi Y, Relman DA, Nishikawa A. Invasion of human respiratory epithelial cells by Bordetella pertussis: possible role for a filamentous hemagglutinin Arg-Gly-Asp sequence and alpha5beta1 integrin. Microb Pathog. 2001;30:279–288.

[51] Bassinet L, Gueirard P, Maitre B, et al. Role of adhesins and toxins in invasion of human tracheal epithelial cells by Bordetella pertussis. Infect Immun. 2000;68:1934–1941.

[52] Stefanelli P, Fazio C, Fedele G, et al. A natural pertactin deficient strain of Bordetella pertussis shows improved entry in human monocyte-derived dendritic cells. New Microbiol. 2009;32:159–166.

[53] Smallridge WE, Rolin OY, Jacobs NT, et al. Different effects of whole-cell and acellular vaccines on Bordetella transmission. J Infect Dis. 2014;209:1981–1988.

